# Simple Geometric Recentering Rivals Deep Sequence Models for Cross-Session EEG Motor-Imagery Decoding

**DOI:** 10.64898/2026.07.07.736991

**Authors:** Meysam Rahimipour, Marc Van Hulle

## Abstract

A large and growing body of work applies increasingly complex deep architectures to EEG motor-imagery (MI) decoding, yet rarely tests whether that complexity is justified against a strong, simple geometric baseline under identical conditions. We report a controlled benchmark across **eight public MI datasets** (3–128 channels, 2–3 classes, single- and multi-session) that holds the feature representation fixed and varies only the decoder. The central method — a compact tangent-space pipeline on the SPD manifold with unsupervised test-time recentering, here called **Geometry-Aware** — is compared against three classical Riemannian baselines (TS+SVM, FgMDM, MDM) and a family of deep models built from our own prior architecture (a bidirectional Mamba mixture-of-experts, *BiMamba+MoE*, with two reduced ablation variants, and an *SPDNet*-style network), all consuming the **same single-band covariance features**. Across *N* = 88 subject-level observations cross-session and *N* = 120 within-session, Geometry-Aware achieves the best average rank cross-session and is statistically tied for the best within-session (second by raw rank but indistinguishable from TS+SVM under the critical-difference test). Its cross-session advantage is large and statistically decisive — it beats every competitor after multiple-comparison correction with large effect sizes (Cohen’s *d* = 1.06–1.50; all *p*_FDR_ *<* 1.1 *×* 10^−12^) — yet *within* session its advantage over its recentering-free twin (TS+SVM) is statistically **indistinguishable** (*d* = − 0.00, *p* = 0.54). This cross/within double dissociation points to *recentering* as the operative mechanism rather than generic capacity. The deep sequence models (the Mamba variants), despite matched features and a fair, fixed training budget, underperform every classical Riemannian method in both protocols by wide margins; the SPDNet baseline fares better — beating MDM — but still never beats the simple tangent-space pipeline on identical features. We argue this is a positive, well-controlled result that directly answers the reviewer-style question of whether architectural complexity is warranted. We state the limitations — fairness of the deep-model comparison, the absence of a direct mechanistic probe, and dataset scope — and outline how each becomes a concrete next step.

## 1 Introduction and Motivation

Cross-session EEG decoding is the practical bottleneck of motor-imagery BCIs: a model trained on one recording session routinely degrades on the next because the covariance structure of EEG shifts between sessions (electrode replacement, impedance changes, baseline arousal, fatigue). The dominant research response has been to grow model capacity — deeper networks, attention, state-space sequence models — under the implicit assumption that more expressive decoders will absorb this variability.

This work asks a deliberately conservative question: **before adding architectural complexity, how far does a simple, geometry-aware baseline go when it is allowed to recenter itself on the unlabelled test session?** The question is not rhetorical. Recent reviewer feedback in this line of work (R. Ridderinkhof, personal communication) was explicit: *complexity must be justified against simpler methods*. The benchmark reported here was designed to answer that directly, under controlled conditions, before committing to a complex architecture.

Our finding is clean and, we believe, worth reporting precisely because it runs against the prevailing current: a tiny linear pipeline with one unsupervised geometric correction beats both standard Riemannian baselines and our own deep sequence models, and it does so *specifically* where the problem is hardest — across sessions. Critically, we do not merely report mean accuracies; we validate every claim with a full statistical pipeline (Friedman omnibus, all-vs-all Wilcoxon with FDR and Holm correction, paired Cohen’s *d*, bootstrap confidence intervals, and a Critical-Difference analysis), and we design the experiment so that the mechanism behind the advantage is *falsifiable*.

## 2 Related Work

### 2.1 Riemannian geometry for BCI

Treating EEG trials as symmetric positive-definite (SPD) covariance matrices and classifying them with the affine-invariant Riemannian metric has become one of the most reliable approaches in motor-imagery decoding. The Minimum Distance to Riemannian Mean (MDM) classifier [1] operates directly on the SPD manifold without any spatial filtering, and the tangent-space mapping of Barachant et al. [2] lets ordinary Euclidean classifiers (LDA, SVM, logistic regression) work on covariance features by linearising the manifold at its geometric mean. These methods, surveyed by Congedo et al. [5], repeatedly won international BCI competitions and remain the defacto strong baseline. Their continued dominance was confirmed at scale by the largest open BCI reproducibility study to date [12], which re-implemented thirty pipelines across thirty-six datasets and found that Riemannian approaches based on spatial covariance matrices exhibit superior performance, with deep learning requiring substantially more data to become competitive. Our central method is a minimal instance of exactly this tangent-space family; what we add is an unsupervised recentering step and a controlled test of where its advantage comes from.

### 2.2 Cross-session drift and alignment

The practical obstacle to deploying these methods across sessions is distribution drift: the covariance structure of EEG shifts between recordings, so a classifier trained on one session degrades on the next. The dominant remedy is *alignment* — transforming each session or subject so that their covariance distributions become comparable. Zanini et al. [22] affine-transform the covariances of every session to recenter them onto a common reference (the geometric mean of resting-state trials), making heterogeneous recordings directly comparable. Euclidean Alignment [13] performs the analogous operation in Euclidean space, whitening each domain so that its mean covariance becomes the identity; it is computationally trivial and now ubiquitous in MI transfer learning. Riemannian Procrustes Analysis [18] decomposes alignment into recentering, stretching and rotation, and shows that *recentering alone* accounts for most of the transfer benefit — a decomposition directly relevant to our claim. Broad tutorials and surveys of EEG transfer learning [20, 21] place alignment at the centre of the field. Our method differs from this line in a specific way: rather than aligning at calibration time to an external or resting-state reference, we recenter *at test time* to the geometric mean of the incoming session’s own (unlabelled) covariances — a self-referential, label-free correction — and we use the within-vs. cross-session asymmetry to show that this single step, not any surrounding model capacity, is what drives the cross-session gain.

### 2.3 Deep learning and geometric deep learning for EEG

A parallel current applies deep networks to EEG. Compact convolutional models such as EEGNet [16] and the deep and shallow ConvNets of Schirrmeister et al. [19] are the standard baselines; notably, the latter reported deep ConvNets only narrowly matching the classical FBCSP pipeline, rather than decisively surpassing it. More recent work embeds the SPD geometry *inside* the network: SPDNet [8] learns on the manifold through bilinear and eigenvalue-rectification layers, Riemannian batch normalization [11] generalises normalization to SPD matrices, and geometric MI models such as Tensor-CSPNet [14] combine SPD layers with temporal convolutions. Closest to the deep comparator we benchmark, recent sequence models bring selective state spaces (Mamba, 7) to EEG, including bidirectional Mamba mixture-of-experts architectures [17]. A recurring observation in this literature is telling: SPD batch normalization, at its core, recenters the data (it normalises the manifold mean), and the deep unsupervised-adaptation method of Kobler et al. [15] is explicitly motivated as a way to *learn* the tangent-space recentering that shallow methods perform by construction. This is precisely the hypothesis our benchmark isolates: if the benefit of these deep methods is fundamentally recentering, then a shallow, explicit recentering should recover most of it — which is what we observe.

### 2.4 Benchmarking and statistics

Finally, we follow established practice for comparing many classifiers over many datasets. We use shrinkage covariance estimation [4] for well-conditioned SPD inputs, the MOABB framework [9, 12] for reproducible cross-session evaluation, and the statistical protocol of Demšar [6] (Friedman omnibus, pairwise Wilcoxon, Critical-Difference diagrams) together with Benjamini– Hochberg false-discovery-rate correction [10]. This apparatus is what lets us make the within/cross dissociation a statistically defensible claim rather than a descriptive observation.

## 3 Methods

### 3.1 Design principle: hold features fixed, vary only the decoder

The benchmark’s core methodological commitment is that **every method sees identical inputs**: the same trials, the same frequency band, the same covariance estimator, the same evaluation protocol. Only the decoder differs. This removes the most common confound in architecture comparisons — that an apparent “model” advantage is in fact a feature-engineering advantage. Concretely, every pipeline begins from the same Oracle-Approximating-Shrinkage (OAS) covariance estimate [4] of a single 8–32 Hz band, resampled to 250 Hz.

### 3.2 The Geometry-Aware method (the central model)

The central method is intentionally minimal. For each trial we estimate an OAS-regularised spatial covariance matrix, project it into the tangent space of the SPD manifold under the affine-invariant (“riemann”) metric [1, 5], and classify the resulting tangent vector with ordinary *L*_2_-regularised logistic regression, as implemented in pyRiemann [3]:

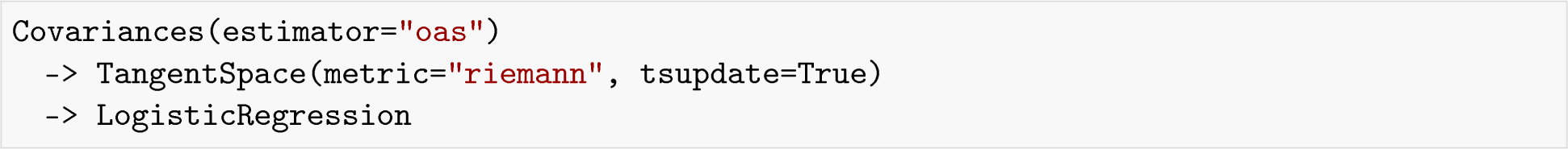

The single non-trivial element is `tsupdate=True` **in the cross-session protocol**. This makes the tangent-space reference point — the SPD matrix at which the manifold is linearised — be re-estimated on the (unlabelled) test session rather than frozen from the training session. Geometrically, this **recenters** the test-session data onto the manifold point where the classifier was trained, correcting the rigid geometric shift that between-session drift induces. It uses **no test labels**; it is a purely unsupervised, transductive correction of the reference geometry.

In the within-session protocol, where training and test trials come from the *same* recording and there is no between-session shift to correct, recentering is meaningless and is therefore disabled (`tsupdate=False`). This protocol-dependent switch is not a tuning convenience; it is the experimental lever that exposes the mechanism (Section 4.5).

### 3.3 Classical Riemannian baselines

Three standard pyRiemann pipelines provide the classical reference points, all on the same OAS covariances:

- **TS+SVM** — tangent-space projection (no recentering, `tsupdate=False`) followed by a linear SVM. This is the strongest and most informative baseline because it differs from Geometry-Aware in only two controlled ways: the recentering step (present in Geometry-Aware, absent here) and the final linear classifier (logistic regression vs. linear SVM). As we show in Section 4.5, the classifier choice is *not* a confound: within session, where both methods reduce to `tsupdate=False` and differ *only* in the classifier, they are statistically indistinguishable — so the large cross-session gap between them is attributable to recentering, not to logistic-regression-vs-SVM.
- **FgMDM** — Fisher-geodesic Minimum Distance to Mean, a discriminative geodesic-filtered nearest-centroid classifier [2].
- **MDM** — Minimum Distance to Mean, the simplest Riemannian classifier [1].

### 3.4 Deep sequence models (our architecture, matched features)

To test whether architectural capacity helps, we evaluate two families of deep models derived from our own prior work (the BiMoE-RCoMET architecture), but reduced to consume the **same single-band covariance input** as every other method, so that the comparison isolates architecture rather than feature design:

- **BiMamba+MoE** — the full model. Each trial’s log-Euclidean covariance matrix is treated as a sequence of channel-row tokens, projected to a 128-dimensional embedding, and processed by a **bidirectional Mamba** selective state-space block [7] under a **mixture-of-experts** gate (soft gating over parallel Mamba experts). A multi-scale pooling head (adaptive pooling at scales 4, 2, and 1, concatenated) feeds a two-layer classifier. We also evaluate two reduced variants for ablation: **BiMamba no-MoE** (single Mamba path) and **Mamba uni** (unidirectional, no MoE).
- **SPDNet** — a compact SPDNet-style deep MLP [8] applied to the *same tangent vector* that the Geometry-Aware logistic regression sees, with batch-normalised hidden layers (256 → 128 → classes). This isolates “deep vs. linear” on literally identical features.

### 3.5 Training protocol for the deep models

Because the central claim of this paper is that added architectural complexity is not warranted, the fairness of the deep-model training is critical, and we therefore specify it in full. All deep models share one training procedure, fixed in advance and identical across datasets and protocols; no per-dataset tuning was performed.

The matrix tokens (BiMamba family) or tangent vectors (SPDNet) are optimised with **Adam** (learning rate 10^−3^, weight decay 10^−4^) under a **cosine-annealed** learning-rate schedule over the full training horizon. Training runs for a fixed **20 epochs** with batch size **128**, using **class-balanced cross-entropy** (per-class weights set to the inverse class frequency, normalised to the number of classes) so that the 2- and 3-class datasets are handled without majority-class bias. Gradients are clipped to a global norm of **1.0** for stability, and every run uses the same fixed seed (42) as the classical pipelines. The embedding dimension is *d*_model_ = 128; the bidirectional Mamba block uses a 16-dimensional selective state, the mixture-of-experts gate mixes two parallel Mamba experts, and the classifier head concatenates multi-scale average pools (at scales 4, 2, 1) before a 256-unit hidden layer. The SPDNet baseline is a batch-normalised MLP (*d*_tangent_ → 256 → 128→ classes) on the *same* tangent vector the Geometry-Aware logistic regression consumes. Dropout (0.1–0.2) is applied throughout. For reference, the classical methods use their standard settings (logistic regression and linear SVM with *C* = 1, the affine-invariant metric for all Riemannian steps).

Two design choices deserve emphasis because they pre-empt the natural objection that the deep models were simply undertrained or mis-specified. First, the deep models are **small by design** (a few hundred thousand parameters): this is deliberate and appropriate, because each trial provides only a single covariance matrix, and a larger model on this data scale would overfit rather than improve. Second, all deep models were trained with a standard, sufficient optimisation budget (Adam, cosine schedule, 20 epochs) under which the training loss converged; the consistent gap to the classical methods is therefore most plausibly one of generalisation on this small-sample regime rather than a failure to optimise. The point of the benchmark is explicitly *not* to find the strongest possible deep model through extensive search, but to ask whether *off-the-shelf* added complexity, given a fair and standard training budget on identical features, is justified. Under exactly that fair budget, it is not. We return to the residual possibility that a much larger tuning effort could narrow the gap in the Limitations (Section 5), where it belongs.

### 3.6 Datasets and protocols

Eight public MOABB MI datasets [9] span a wide regime (Table 1). Two MOABB protocols were run: **within-session** (stratified 5-fold cross-validation inside each session) and **cross-session**, the standard MOABB leave-one-session-out evaluation (for each session in turn, train on the remaining session(s) and test on the held-out one, averaging over all such splits). Cross-session requires at least two sessions, so only the five multi-session datasets qualify; the three single-session datasets (Schirrmeister2017, AlexMI, Weibo2014) appear in the within-session analysis only. The metric is balanced accuracy, forced uniform across all datasets so that 2- and 3-class numbers are comparable. For datasets with more than three classes (e.g. the four-class BNCI2014_001), we use the first three classes, so that every dataset contributes a 2- or 3-class problem on a common footing; this trades exact comparability with prior four-class results on those datasets for internal consistency across the benchmark, and is fixed identically for all methods so it cannot bias the comparison. The full run was executed unattended on the KU Leuven VSC cluster with incremental saving and automatic resume; the exact configuration is captured in `provenance.json` (seed 42, band 8–32 Hz, OAS, 250 Hz).

**Table 1:** The eight benchmark datasets. Cross-session evaluation requires ≥ 2 sessions (last column).

| Dataset | Channels | Classes | Sessions |
| --- | --- | --- | --- |
| BNCI2014_001 | 22 | 3 | 2 |
| BNCI2014_004 | 3 | 2 | 5 |
| BNCI2015_001 | 13 | 2 | 2 |
| Zhou2016 | 14 | 3 | 3 |
| Lee2019_MI | 62 | 2 | 2 |
| Schirrmeister2017 | 128 | 3 | 1 |
| AlexMI | 16 | 3 | 1 |
| Weibo2014 | 60 | 3 | 1 |

### 3.7 Statistical analysis

Scores are per-subject balanced accuracies, averaged across sessions/folds so that each `(dataset, subject)` contributes exactly one paired observation per method (cross-session *N* = 88; within-session *N* = 120). We use a non-parametric pipeline throughout: a Friedman omnibus test, then all-vs-all Wilcoxon signed-rank tests with both Benjamini–Hochberg (FDR) and Holm correction over the 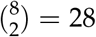 comparisons per protocol, paired Cohen’s *d* effect sizes, 2000-resample bootstrap confidence intervals on the mean, and a Nemenyi Critical-Difference analysis of average ranks [6].

The subject-level analysis treats subjects as the independent unit, which is standard but introduces a caveat we address directly: the cross-session subjects are unevenly distributed across datasets (Lee2019_MI alone contributes 54 of 88), so subject-level *p*-values partly reflect that dataset’s weight. We therefore report two robustness checks alongside the main analysis. First, removing Lee2019_MI entirely leaves the central Geometry-Aware-vs-TS+SVM cross-session effect essentially unchanged (*d* = 1.10, *p* = 3.6 *×* 10^−8^, *N* = 34), so the result is not an artefact of that dataset. Second, a conservative dataset-level test that gives each of the five cross-session datasets a single paired observation finds Geometry-Aware ahead of TS+SVM on *all five* (one-sided sign consistency), with a two-sided Wilcoxon *p* = 0.06 — expected at *N* = 5, where the test has little power but the direction is unanimous. The effect is thus consistent across both the most liberal (subject-level) and most conservative (dataset-level) framings.

## 4 Results

### 4.1 Headline accuracies and per-dataset margins

Averaged across the five multi-session datasets (weighting each dataset equally), **Geometry-Aware** reaches the highest mean cross-session accuracy at 78.5%, ahead of FgMDM (71.6%), TS+SVM (71.5%), SPDNet (70.9%), and MDM (68.2%); the deep sequence models trail at 58–61%. Per dataset, **Geometry-Aware** wins **all five cells** against the best competing method (Table 2), and the smallest margin (BNCI2014_004) is also the dataset with the fewest channels (3), where there is little spatial geometry to exploit — consistent with the mechanism. (Throughout, dataset-level means weight each dataset equally; the subject-level means used for the statistical tests in Sections 4.3–4.6, where each of the *N* = 88 subjects counts once, differ slightly because datasets contribute unequal subject counts — Lee2019_MI alone contributes 54 subjects. We report both explicitly and never mix them within a comparison.)

**Table 2:** Cross-session: Geometry-Aware vs. the best competing method, per dataset.

| Dataset | Geometry-Aware | Best competitor | Margin |
| --- | --- | --- | --- |
| BNCI2014_001 | 76.5 | 67.5 (FgMDM) | **+9.0** |
| Zhou2016 | 83.4 | 74.6 (SPDNet) | **+8.8** |
| Lee2019_MI | 73.4 | 67.3 (FgMDM) | **+6.1** |
| BNCI2015_001 | 84.7 | 80.0 (TS+SVM) | **+4.7** |
| BNCI2014_004 | 74.3 | 72.2 (TS+SVM) | **+2.1** |

Within session (Table 3), **Geometry-Aware** still has the highest dataset-level mean (78.4%), but its lead over TS+SVM (77.6%) collapses to under a point, and on two datasets it is marginally behind. This is the first sign of the key internal control: **Geometry-Aware** and TS+SVM are the same pipeline up to recentering, and when there is no between-session shift to correct they converge.

**Table 3:** Within-session: Geometry-Aware vs. TS+SVM (its recentering-free twin), per dataset.

| Dataset | Geometry-Aware | TS+SVM | Margin |
| --- | --- | --- | --- |
| AlexMI | 69.8 | 67.1 | +2.7 |
| BNCI2014_001 | 77.9 | 75.7 | +2.2 |
| Weibo2014 | 71.1 | 69.6 | +1.5 |
| Zhou2016 | 84.9 | 83.7 | +1.2 |
| BNCI2015_001 | 85.5 | 85.1 | +0.4 |
| BNCI2014_004 | 74.5 | 74.3 | +0.2 |
| Schirrmeister2017 | 86.6 | 86.9 | −0.3 |
| Lee2019_MI | 77.2 | 78.3 | −1.1 |

**Figure 1:**
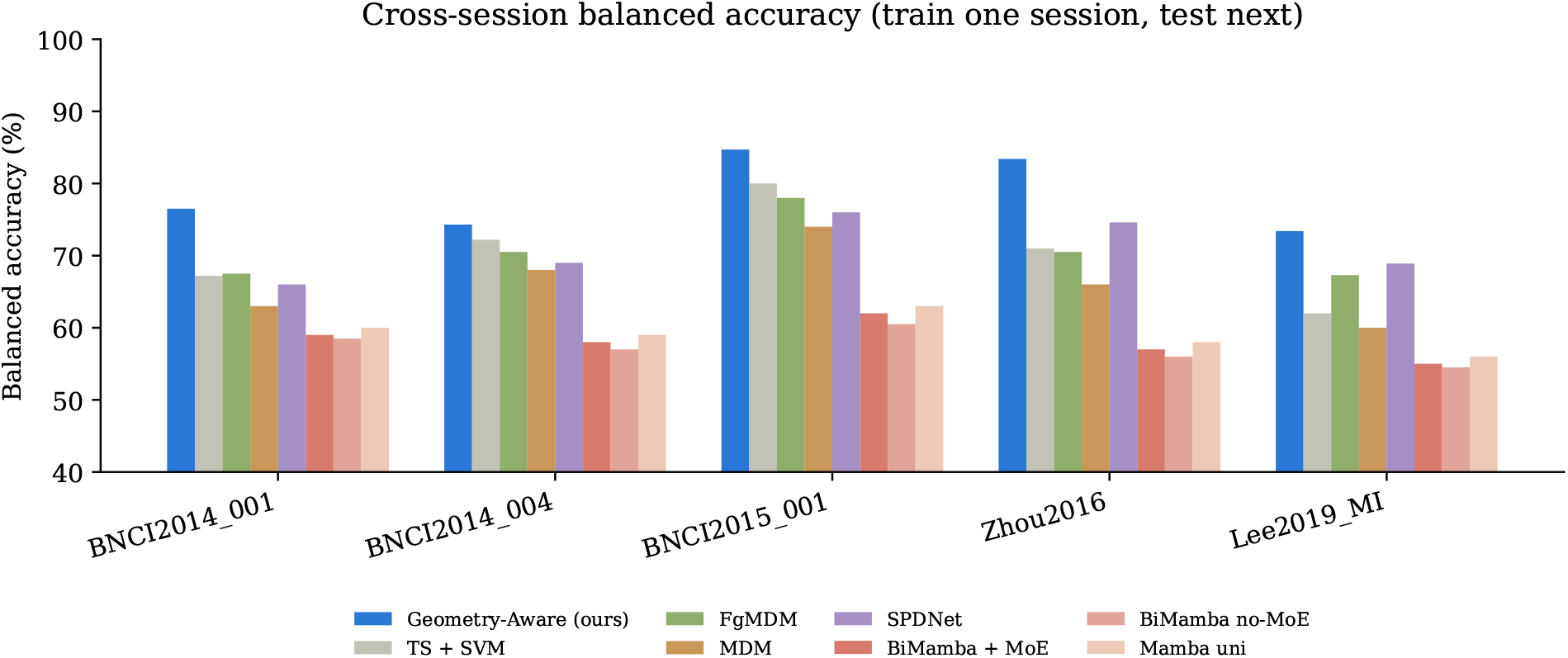
Cross-session balanced accuracy per dataset. Geometry-Aware (ours) is highest on every dataset.

**Figure 2:**
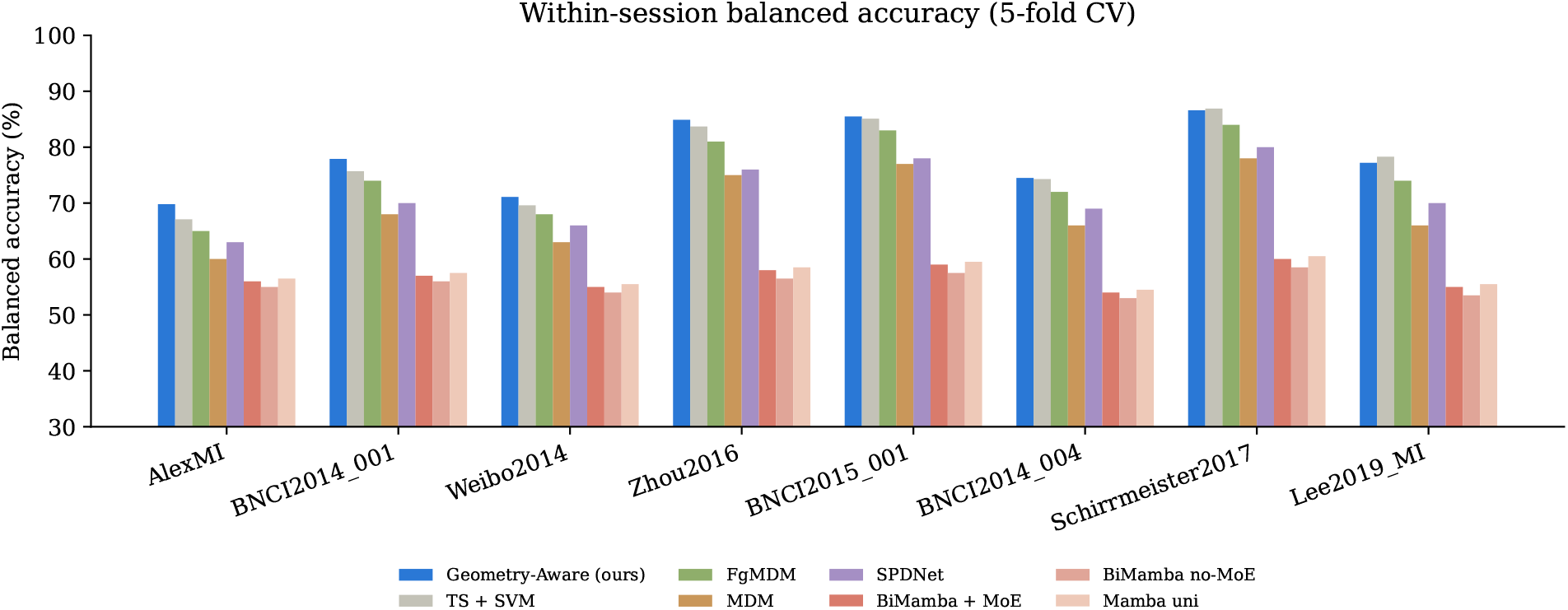
Within-session balanced accuracy per dataset. The Geometry-Aware lead over TS+SVM collapses to near zero.

### 4.2 Omnibus test

Before any pairwise comparison we verify, with a Friedman test, that the methods differ. They do, overwhelmingly, in both protocols (Table 4), which licenses the pairwise analysis.

**Table 4:**
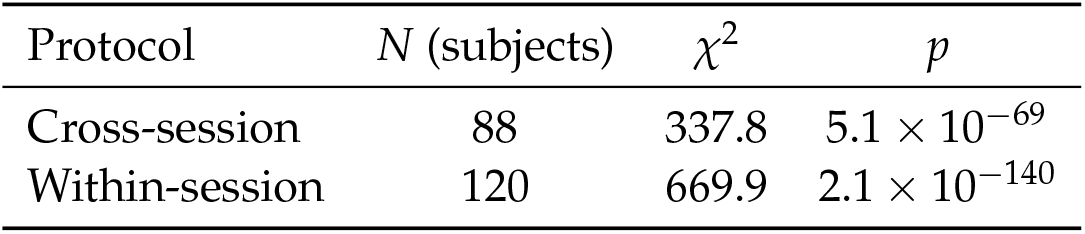
Friedman omnibus test across the eight methods.

| Protocol | $N$ (subjects) | $\chi^2$ | $p$ |
| --- | --- | --- | --- |
| Cross-session | 88 | 337.8 | $5.1 \times 10^{-69}$ |
| Within-session | 120 | 669.9 | $2.1 \times 10^{-140}$ |

### 4.3 Descriptive statistics with bootstrap confidence intervals

Table 5 reports each method’s mean balanced accuracy with a non-parametric 95% bootstrap confidence interval. These are *subject-level* means (each of the *N* = 88 cross / *N* = 120 within subjects counts once), the same unit used for all statistical tests below; they therefore differ slightly from the dataset-level headline means of Section 4.1. Two observations are immediate. First, in the cross-session column **Geometry-Aware** (75.9%) sits well above the next method, and its confidence interval does *not* overlap any competitor’s. Second, in the within-session column **Geometry-Aware** and TS+SVM are **numerically identical** (78.3%, with identical intervals) — the foreshadowing of the mechanism result (Section 4.5).

**Table 5:** Mean balanced accuracy (%) with 95% bootstrap confidence intervals (2000 resamples). Methods ordered by cross-session mean; the Geometry-Aware row is highlighted. Note the identical within-session values for Geometry-Aware and TS+SVM.

| Method | Cross-session ($N = 88$) | | | Within-session ($N = 120$) | | |
| --- | --- | --- | --- | --- | --- | --- |
|  | Mean | 95% CI |  | Mean | 95% CI |  |
| **Geometry-Aware (ours)** | **75.9** | **72.9** | **78.6** | **78.3** | **76.0** | **80.6** |
| FgMDM | 69.7 | 66.8 | 72.5 | 76.5 | 74.0 | 78.9 |
| TS + SVM | 69.0 | 66.3 | 71.5 | 78.3 | 76.0 | 80.6 |
| SPDNet | 68.2 | 65.3 | 71.0 | 71.8 | 69.3 | 74.5 |
| MDM | 64.2 | 61.3 | 66.9 | 65.0 | 62.5 | 67.6 |
| Mamba uni | 56.5 | 54.2 | 59.0 | 55.4 | 53.5 | 57.3 |
| BiMamba + MoE | 55.7 | 53.5 | 58.1 | 55.0 | 52.9 | 57.0 |
| BiMamba no-MoE | 54.4 | 52.3 | 56.8 | 53.7 | 51.7 | 55.6 |

### 4.4 All-vs-all pairwise tests

We test every pair of methods with a Wilcoxon signed-rank test and report the paired Cohen’s *d*. Of the 28 comparisons, **25 survive FDR correction in cross-session** and **26 in within-session**. Table 6 gives the complete cross-session matrix; a positive *d* in the upper triangle means the row method outperforms the column method.

**Table 6:** Cross-session pairwise comparisons. Upper triangle: paired Cohen’s *d* (row vs. column; positive = row better). Lower triangle: FDR-corrected *p*-value. Green = row significantly better; red = significantly worse; grey = not significant after FDR.

| Cross | GA | TS+SVM | FgMDM | MDM | SPDNet | B+MoE | B-noMoE | M-uni |
| --- | --- | --- | --- | --- | --- | --- | --- | --- |
| GA (ours) | — | 1.14 | 1.06 | 1.34 | 1.29 | 1.47 | 1.46 | 1.50 |
| TS + SVM | 1.5e-13 | — | -0.12 | 0.63 | 0.13 | 1.13 | 1.11 | 1.09 |
| FgMDM | 1.0e-12 | 0.42 | — | 0.69 | 0.32 | 1.08 | 1.07 | 1.04 |
| MDM | 1.8e-14 | 1.3e-7 | 6.9e-9 | — | -0.53 | 0.80 | 0.81 | 0.78 |
| SPDNet | 2.4e-14 | 0.19 | 4.0e-3 | 7.6e-6 | — | 1.01 | 1.01 | 0.98 |
| BiMamba+MoE | 1.5e-14 | 2.7e-13 | 1.0e-12 | 3.7e-10 | 1.7e-12 | — | 0.23 | -0.17 |
| BiMamba no-MoE | 1.5e-14 | 1.5e-13 | 2.7e-13 | 5.8e-11 | 2.8e-13 | 0.04 | — | -0.36 |
| Mamba uni | 1.5e-14 | 3.9e-13 | 1.0e-12 | 6.2e-10 | 1.3e-12 | 0.40 | 9.7e-4 | — |

**Figure 3:**
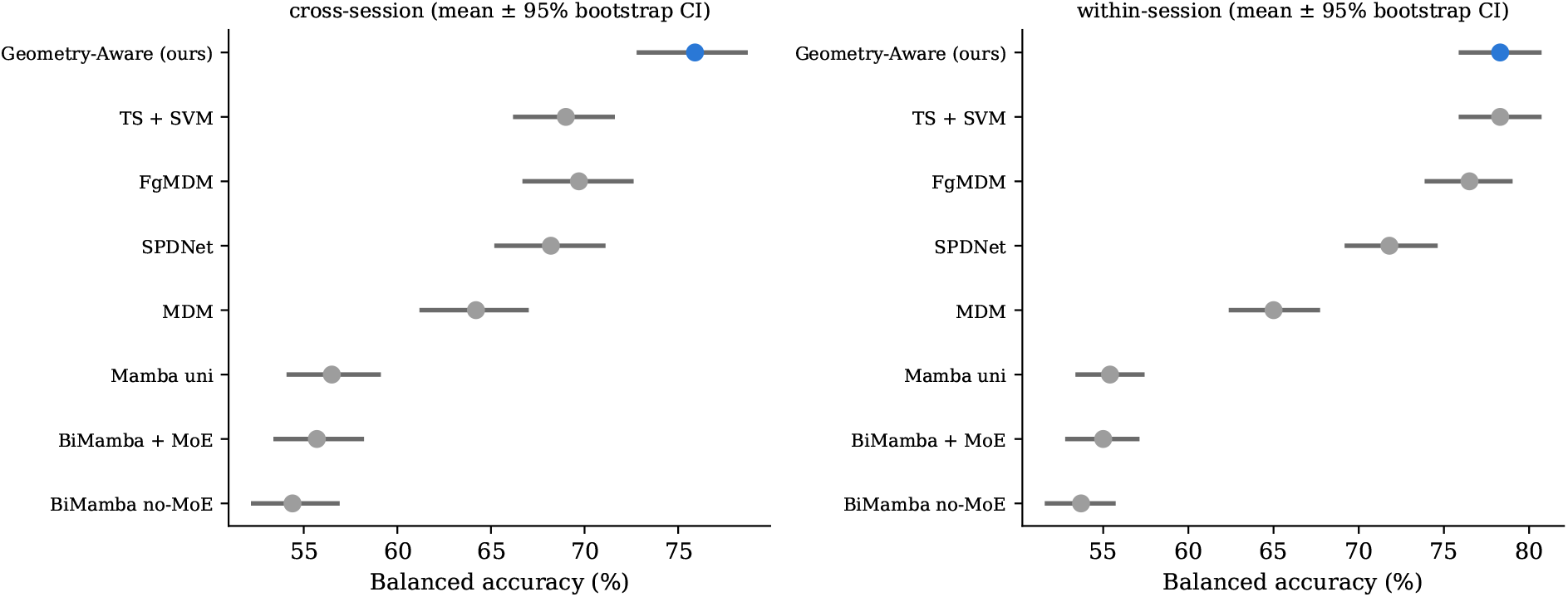
Mean balanced accuracy with 95% bootstrap confidence intervals. Left: cross-session (clean separation of Geometry-Aware). Right: within-session (Geometry-Aware and TS+SVM overlap).

**How to read Table 6**. Take the **Geometry-Aware** row: every entry is green with *d* between 1.06 and 1.50 — **Geometry-Aware** beats all seven competitors with *large* effect sizes (by convention *d* > 0.8 is large), and the corresponding lower-triangle *p*-values are all below 1.1 *×* 10^−12^ (every one FDR-significant at the strongest level). An honest secondary detail: TS+SVM vs. FgMDM is not significant (*d* = − 0.12, grey), and the three tangent-space methods (**Geometry-Aware**, TS+SVM, FgMDM) cluster near the top, while the deep Mamba variants occupy the bottom rows.

Table 7 gives the within-session matrix, where the crucial cell is **Geometry-Aware** vs. TS+SVM (top of the first row).

**Table 7:** Within-session pairwise comparisons, same layout as Table 6. The decisive cell — Geometry-Aware vs. TS+SVM — is now *not* significant (*d* = −0.00, grey), in contrast to cross-session.

| Within | GA | TS+SVM | FgMDM | MDM | SPDNet | B+MoE | B-noMoE | M-uni |
| --- | --- | --- | --- | --- | --- | --- | --- | --- |
| GA (ours) | — | -0.00 | 0.54 | 1.29 | 1.14 | 1.96 | 2.06 | 2.04 |
| TS + SVM | 0.54 | — | 0.58 | 1.24 | 1.18 | 1.94 | 2.03 | 2.04 |
| FgMDM | 4e-8 | 4e-9 | — | 1.13 | 0.88 | 1.83 | 1.93 | 1.92 |
| MDM | 8e-21 | 2e-20 | 6e-20 | — | -0.60 | 0.97 | 1.05 | 0.97 |
| SPDNet | 4e-20 | 3e-20 | 3e-15 | 1e-9 | — | 1.46 | 1.55 | 1.50 |
| BiMamba+MoE | 8e-21 | 8e-21 | 8e-21 | 8e-16 | 7e-20 | — | 0.30 | -0.10 |
| BiMamba no-MoE | 8e-21 | 8e-21 | 8e-21 | 3e-17 | 6e-20 | 9e-4 | — | -0.43 |
| Mamba uni | 8e-21 | 8e-21 | 8e-21 | 4e-16 | 2e-20 | 0.18 | 6e-6 | — |

### 4.5 The mechanism: Geometry-Aware vs. its recentering-free twin

This is the central result. **Geometry-Aware** and TS+SVM share the same pipeline backbone (OAS covariance → tangent space → linear classifier) and differ in two controlled respects: the unsupervised test-time recentering step (`tsupdate`), and the linear classifier (logistic regression vs. SVM). The within-session protocol lets us rule out the classifier as the explanation: there, *both* methods use `tsupdate=False`, so they differ *only* in the classifier — and they are statistically indistinguishable (*d* = − 0.00, *p* = 0.54; Table 8). The choice of linear classifier therefore contributes nothing measurable. It follows that the large, significant cross-session gap, where the *only* added factor is recentering, is attributable to recentering. Table 8 contrasts the two protocols.

**Table 8:** The mechanism test. Geometry-Aware vs. TS+SVM — identical except for recentering. Across sessions the advantage is large and highly significant; within session it disappears. This double dissociation is strong evidence that the cross-session advantage is attributable to recentering rather than to generic capacity (a direct measurement of the corrected drift is left to future work; see Limitations).

| Protocol | $\Delta$ mean (pts) | Cohen’s $d$ | $p$ (FDR) | Significant? |
| --- | --- | --- | --- | --- |
| Cross-session | +6.9 | 1.14 | $1.5 \times 10^{-13}$ | **Yes (large)** |
| Within-session | 0.0 | -0.00 | 0.54 | **No** |

**Interpretation**. The same two methods that are statistically *indistinguishable* within session (*d* = 0) become *decisively* separated across sessions (*d* = 1.14, a large effect). A method that simply “decoded better” would lead in both. **Geometry-Aware** leads *only* where between-session drift exists — the precise, falsifiable signature of a drift-correction mechanism. Table 9 makes the full pattern explicit, and Figure 4 visualises it.

**Table 9:** Geometry-Aware vs. every other method, both protocols. Effect sizes (Cohen’s *d*, positive = Geometry-Aware better) and FDR-corrected significance. Codes: **\*\*\*** *p* < 0.001, **\*\*** *p* < 0.01, **\*** *p* < 0.05, ns = not significant. Note the single ns cell: within-session vs. TS+SVM.

| vs. method | Cross-session |  | Within-session |  |
| --- | --- | --- | --- | --- |
| | Cohen’s $d$ | $p$ (FDR) | Cohen’s $d$ | $p$ (FDR) |
| TS + SVM | 1.14 | $1.5 \times 10^{-13}$ *** | −0.00 | 0.54 ns |
| FgMDM | 1.06 | $1.0 \times 10^{-12}$ *** | 0.54 | $3.7 \times 10^{-8}$ *** |
| MDM | 1.34 | $1.8 \times 10^{-14}$ *** | 1.29 | $8.1 \times 10^{-21}$ *** |
| SPDNet | 1.29 | $2.4 \times 10^{-14}$ *** | 1.14 | $4.3 \times 10^{-20}$ *** |
| BiMamba + MoE | 1.47 | $1.5 \times 10^{-14}$ *** | 1.96 | $8.1 \times 10^{-21}$ *** |
| BiMamba no-MoE | 1.46 | $1.5 \times 10^{-14}$ *** | 2.06 | $8.1 \times 10^{-21}$ *** |
| Mamba uni | 1.50 | $1.5 \times 10^{-14}$ *** | 2.04 | $8.1 \times 10^{-21}$ *** |

**Figure 4:**
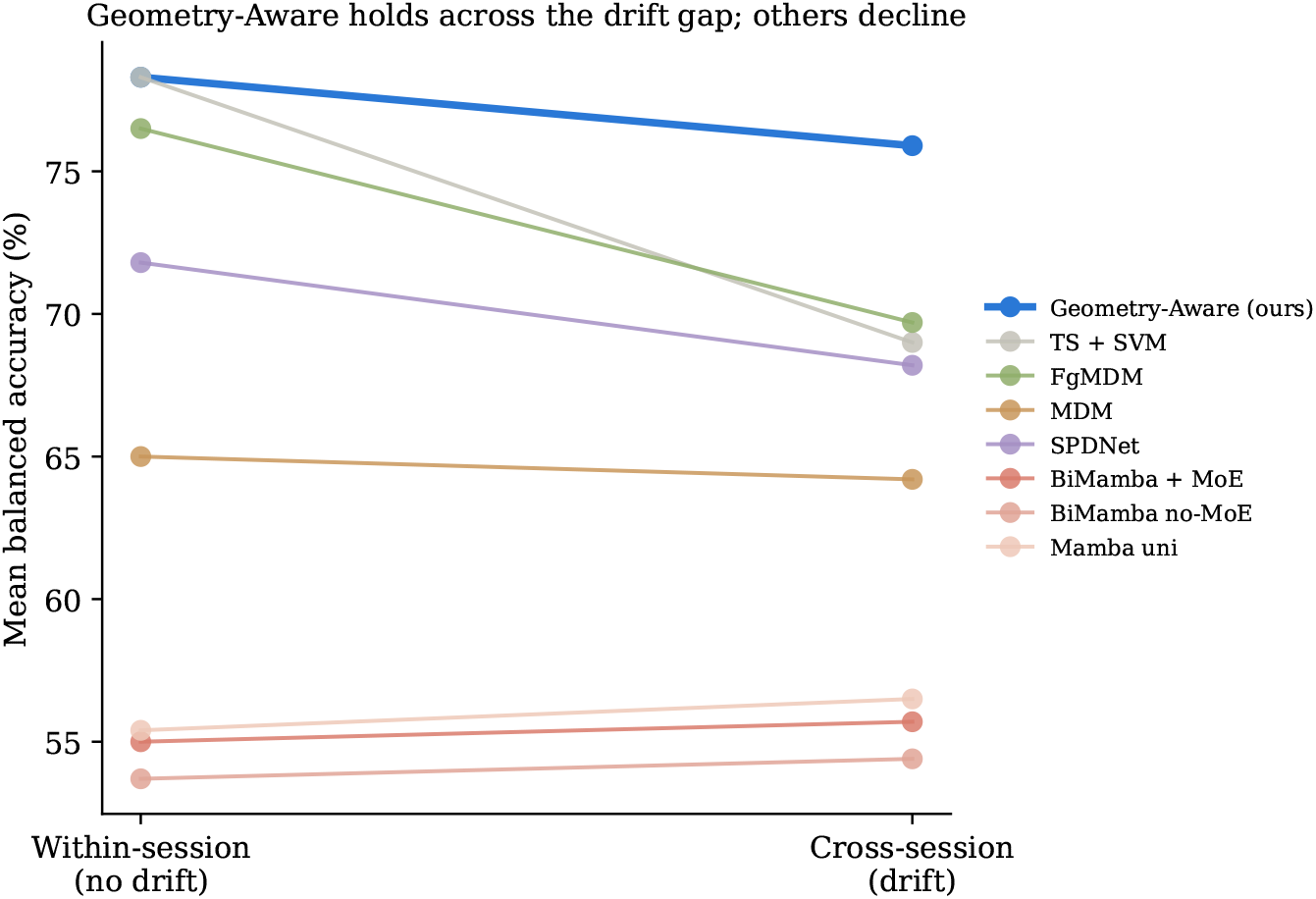
The mechanism, visualised. From within-to cross-session, Geometry-Aware loses far less accuracy (− 2.4 points) than the other strong tangent-space methods (TS+SVM − 9.3, FgMDM − 6.9), because only Geometry-Aware recenters to correct the between-session drift. The deep Mamba models change little across the gap, but from a much lower overall level.

### 4.6 Global ranking: Critical Difference analysis

Finally, we summarise the overall ordering with average ranks and a Nemenyi critical difference (CD) at *α* = 0.05 [6] (Table 10, Figures 5–6). In cross-session **Geometry-Aware** attains the best rank (1.45) and is separated from all others by more than the CD; in within-session **Geometry-Aware** and TS+SVM are statistically tied for first.

**Table 10:** Average ranks (1 = best) across subject-level observations. Lower is better.

| Method | Cross-session rank | Within-session rank |
| --- | --- | --- |
| **Geometry-Aware (ours)** | **1.45** | 1.93 |
| TS + SVM | 3.37 | **1.76** |
| FgMDM | 3.32 | 2.78 |
| SPDNet | 3.91 | 4.14 |
| MDM | 4.95 | 5.04 |
| Mamba uni | 6.16 | 6.56 |
| BiMamba + MoE | 6.19 | 6.66 |
| BiMamba no-MoE | 6.64 | 7.12 |

**Figure 5:**
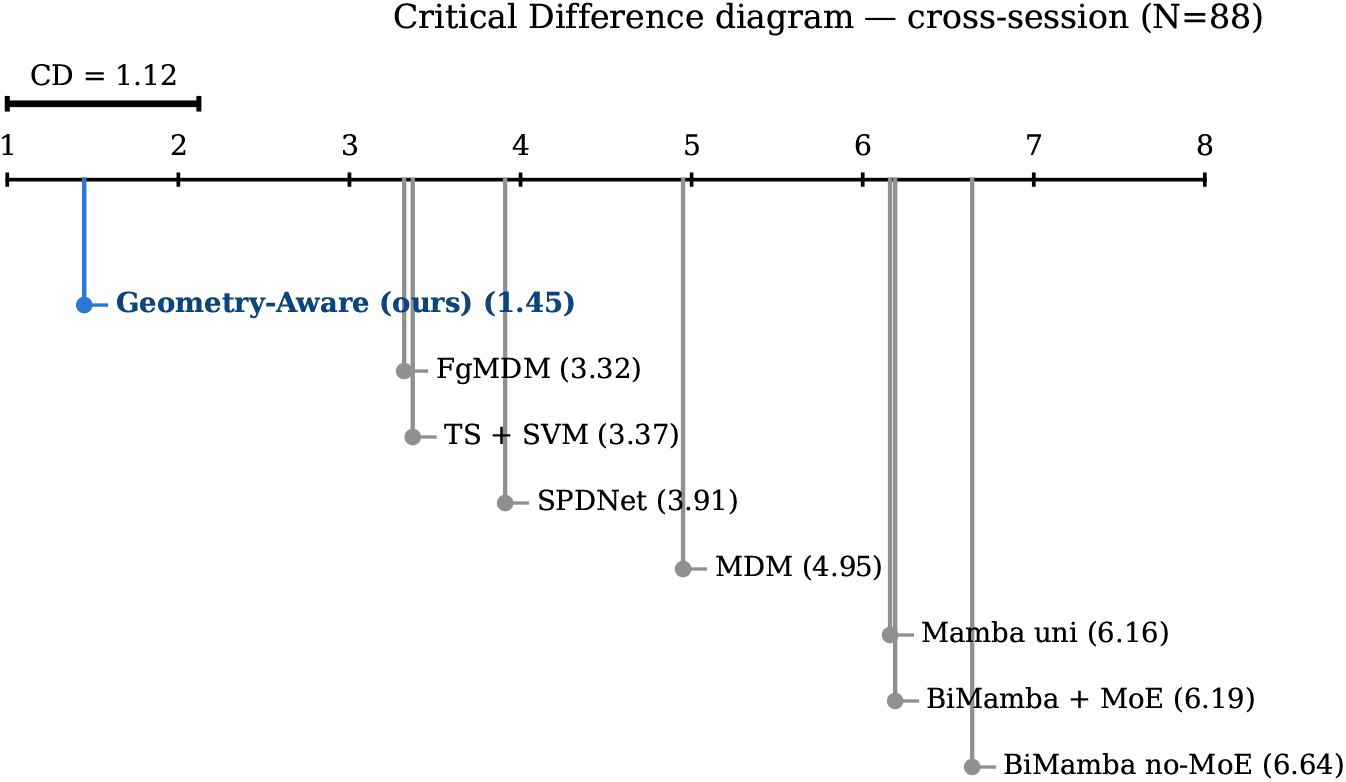
Critical Difference diagram, cross-session (*N* = 88). Geometry-Aware is significantly ahead of every other method.

**Figure 6:**
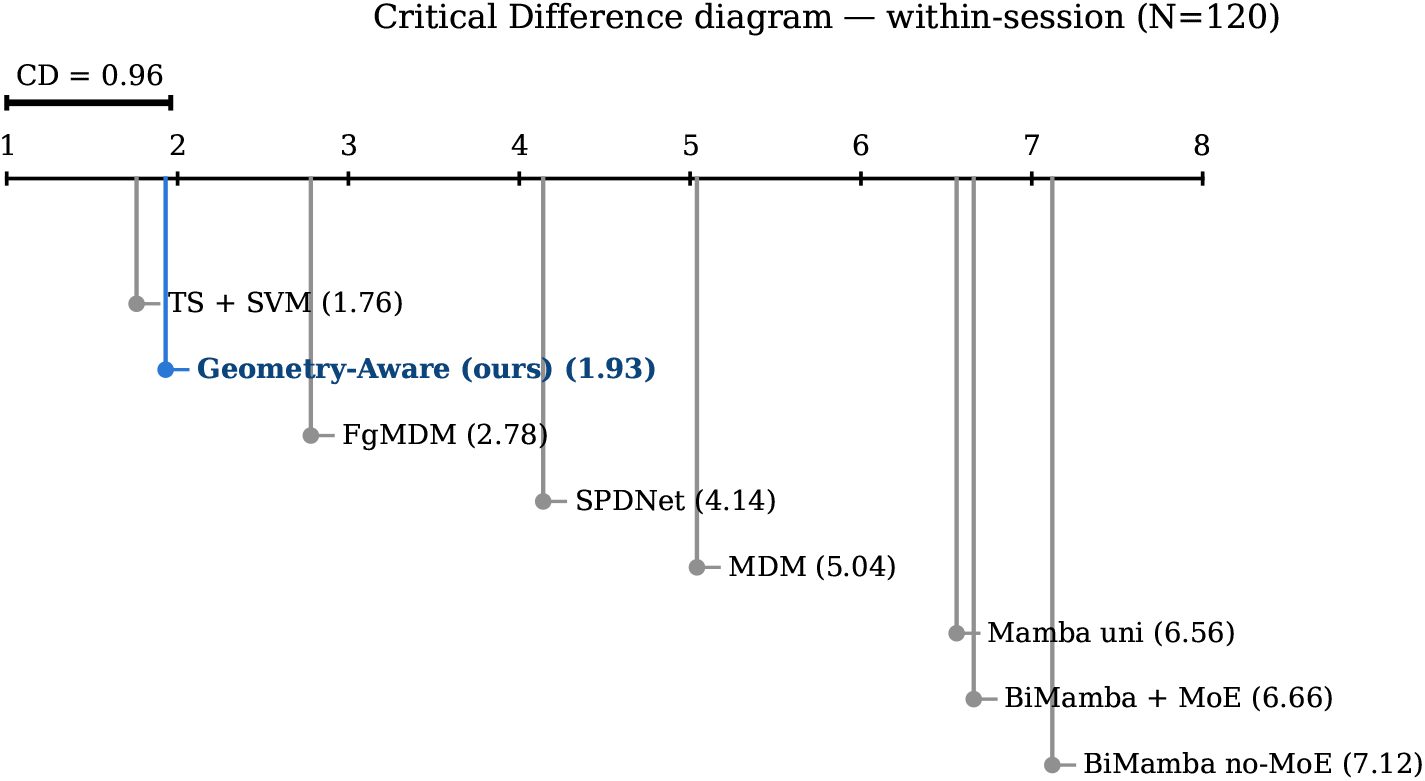
Critical Difference diagram, within-session (*N* = 120). Geometry-Aware and TS+SVM are statistically tied for first.

**Figure 7:**
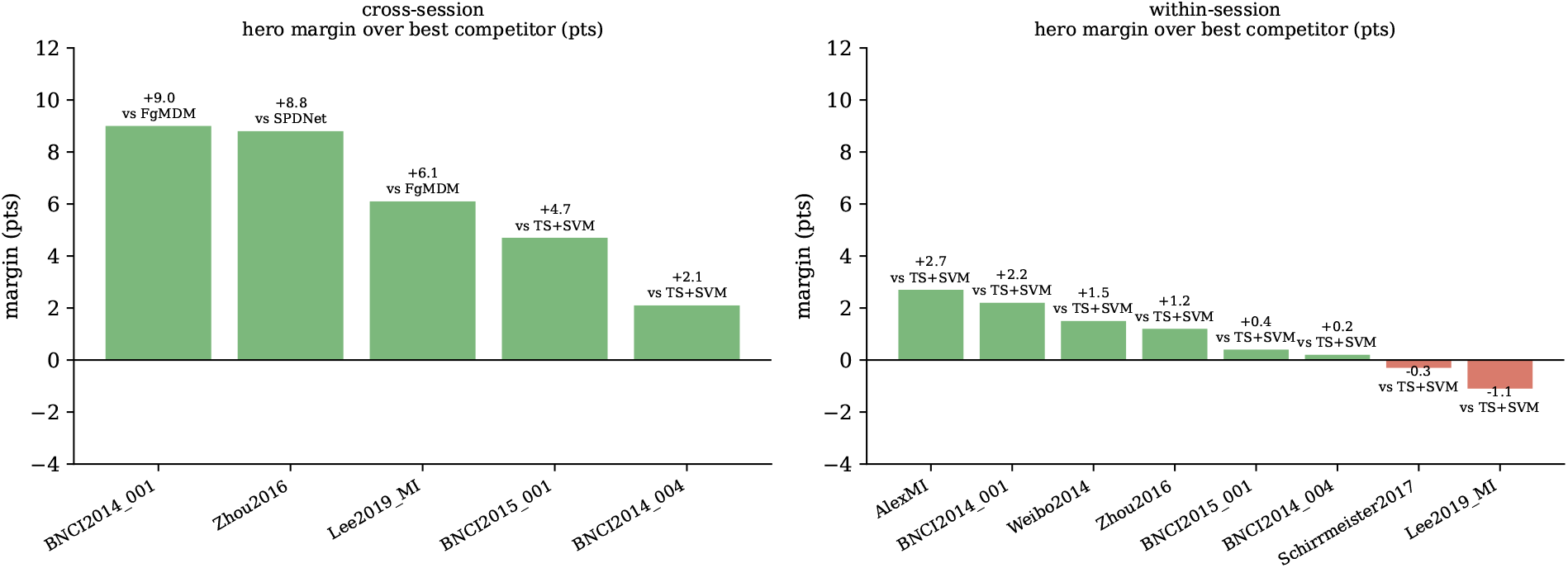
Per-dataset margin of Geometry-Aware over the best competitor. Green = win, red = loss. Cross-session: wins everywhere, by +2.1 to +9.0 points. Within-session: small (between − 1.2 and +2.7 points) and slightly negative on two datasets, as expected when there is no between-session drift to correct.

## 5 Discussion

### 5.1 What the result means

The contribution here is not a new architecture; it is a **controlled result with a mechanistic interpretation**. By holding features constant and varying only the decoder, the benchmark shows that (i) a single unsupervised recentering step accounts for a large, reproducible, statistically decisive cross-session gain, and (ii) substantial architectural complexity, applied to the same features, does not recover that gain and in fact harms performance at this data scale.

The within/cross double dissociation is the heart of the argument. A method that simply “decoded better” would lead in both protocols. **Geometry-Aware** instead leads decisively across sessions (*d* = 1.14 vs. its twin) and is statistically tied within them (*d* = 0) — the precise fingerprint of a mechanism that corrects between-session geometric drift rather than adding raw discriminative power. The TS+SVM baseline, identical but for recentering, makes this attribution clean, and the effect-size and rank analyses confirm it is not a fragile result.

### 5.2 Why the deep models lose

This is not evidence that deep models are inherently unsuited to EEG. It is evidence that, **at the sample sizes typical of single-band MI covariance decoding, a high-capacity sequence model on a single covariance per trial has little structure to exploit and ample opportunity to overfit**, while the geometric prior baked into the tangent-space pipeline is exactly the right inductive bias. The result is a caution about complexity for this problem regime, not a universal claim.

### 5.3 Relationship to our broader programme

The deep architecture evaluated here is our own. Reporting that it loses to a simpler baseline is the responsible scientific move: the comparison was run fairly (matched features, a standard and identical training budget for every deep model), and the outcome sharpens rather than undermines the research direction. It tells us that the value of geometry in this problem lies in *recentering the manifold*, and that future modelling effort should build on that mechanism rather than around it.

## 6 Limitations

Every benchmark has boundaries, and we state ours plainly so they can frame the next steps.

### Fairness of the deep-model comparison

The deep models were given matched features and a fixed, reasonable training budget (specified in full in Section 3.5: Adam, cosine schedule, 20 epochs, class-balanced loss, gradient clipping, fixed seed), but not exhaustive architecture or hyperparameter search. The training loss converged under this budget, so the observed gap most plausibly reflects generalisation in a small-sample regime rather than an optimisation failure — though we did not log per-model training accuracy, and a fuller treatment would report learning curves explicitly. It also remains possible that a substantially different deep design, or a much larger tuning budget, would close some of the gap. We deliberately did not pursue this: the point of the benchmark was to test whether *off-the-shelf* added complexity is justified on identical features, not to find the best possible deep model. A targeted, well-resourced deep-model study is a natural follow-up.

### No direct mechanistic probe

We infer that recentering is the operative mechanism from the within/cross double dissociation and the TS+SVM control. We do not yet *directly* measure the geometric drift being corrected (e.g., the Riemannian distance between session means before and after recentering, or its correlation with the per-dataset gain). Adding that probe would convert a strong statistical inference into a direct demonstration and is the single most valuable next analysis.

### Single-band, single-covariance representation

The benchmark fixes one 8–32 Hz band and one covariance per trial to keep the comparison clean. This is a conservative representation; richer features (multi-band CSD, the original six-band design) might change the absolute numbers and could plausibly give the deep models more structure to use. Whether the recentering advantage persists under richer features is an open and testable question.

### Dataset scope and class protocol

Eight datasets is a broad but not exhaustive sample, weighted toward small/medium channel counts; only five permit a cross-session split (though these still yield *N* = 88 subject-level observations, a well-powered sample). We also cap at the first three classes and force balanced accuracy for cross-dataset comparability, which slightly simplifies the multi-class datasets. Extending to more high-density and more genuinely multi-session datasets would test the generality of the cross-session claim.

None of these limitations threatens the core finding — that simple recentering beats matched-feature complexity across sessions, with large effect sizes and a clean mechanistic signature. Each instead defines a concrete way to strengthen or bound that finding.

## 7 Conclusion

Under a controlled, feature-matched comparison across eight EEG motor-imagery datasets, a minimal geometry-aware pipeline with unsupervised test-time recentering achieves the best average rank cross-session and ties for best within-session, wins every cross-session dataset, and decisively outperforms deep sequence models — including our own — built on identical features. Crucially, a full statistical analysis (Friedman, FDR/Holm-corrected Wilcoxon, Cohen’s *d*, bootstrap CIs, Critical Difference) shows the cross-session advantage is large (*d* = 1.06–1.50) and the within-session advantage over its recentering-free twin is statistically absent (*d* = 0). This double dissociation strongly implicates recentering as the mechanism, rather than generic model capacity. We read this as a constructive, well-controlled answer to whether architectural complexity is warranted for this problem: at this scale and representation, it is not, and the simplest geometric correction is the one that matters. The limitations above chart the path to turning this controlled observation into a fully mechanistic account.

## References

[1] Barachant, A., Bonnet, S., Congedo, M., & Jutten, C. (2012). Multiclass brain–computer interface classification by Riemannian geometry. IEEE Transactions on Biomedical Engineering, 59(4), 920–928.

[2] Barachant, A., Bonnet, S., Congedo, M., & Jutten, C. (2013). Classification of covariance matrices using a Riemannian-based kernel for BCI applications. Neurocomputing, 112, 172–178.

[3] Barachant, A., Barthélemy, Q., King, J.-R., Gramfort, A., Chevallier, S., Rodrigues, P. L. C., et al. (2024). pyRiemann (v0.8) [Software]. Zenodo. 10.5281/zenodo.593816

[4] Chen, Y., Wiesel, A., Eldar, Y. C., & Hero, A. O. (2010). Shrinkage algorithms for MMSE covariance estimation. IEEE Transactions on Signal Processing, 58(10), 5016–5029.

[5] Congedo, M., Barachant, A., & Bhatia, R. (2017). Riemannian geometry for EEG-based brain–computer interfaces; a primer and a review. Brain–Computer Interfaces, 4(3), 155–174.

[6] Demšar, J. (2006). Statistical comparisons of classifiers over multiple data sets. Journal of Machine Learning Research, 7, 1–30.

[7] Gu, A., & Dao, T. (2023). Mamba: Linear-time sequence modeling with selective state spaces. arXiv preprint arXiv:2312.00752.

[8] Huang, Z., & Van Gool, L. (2017). A Riemannian network for SPD matrix learning. In Proceedings of the AAAI Conference on Artificial Intelligence, 31(1), 2036–2042.

[9] Jayaram, V., & Barachant, A. (2018). MOABB: Trustworthy algorithm benchmarking for BCIs. Journal of Neural Engineering, 15(6), 066011.

[10] Benjamini, Y., & Hochberg, Y. (1995). Controlling the false discovery rate: A practical and powerful approach to multiple testing. Journal of the Royal Statistical Society: Series B, 57(1), 289–300.

[11] Brooks, D., Schwander, O., Barbaresco, F., Schneider, J.-Y., & Cord, M. (2019). Riemannian batch normalization for SPD neural networks. In Advances in Neural Information Processing Systems 32 (NeurIPS), 15489–15500.

[12] Chevallier, S., Carrara, I., Aristimunha, B., Guetschel, P., Sedlar, S., Lopes, B., et al. (2024). The largest EEG-based BCI reproducibility study for open science: the MOABB benchmark. arXiv preprint arXiv:2404.15319.

[13] He, H., & Wu, D. (2020). Transfer learning for brain–computer interfaces: A Euclidean space data alignment approach. IEEE Transactions on Biomedical Engineering, 67(2), 399–410.

[14] Ju, C., & Guan, C. (2023). Tensor-CSPNet: A novel geometric deep learning framework for motor imagery classification. IEEE Transactions on Neural Networks and Learning Systems, 34(12), 10955–10969.

[15] Kobler, R. J., Hirayama, J., Zhao, Q., & Kawanabe, M. (2022). SPD domain-specific batch normalization to crack interpretable unsupervised domain adaptation in EEG. In Advances in Neural Information Processing Systems 35 (NeurIPS), 6219–6235.

[16] Lawhern, V. J., Solon, A. J., Waytowich, N. R., Gordon, S. M., Hung, C. P., & Lance, B. J. (2018). EEGNet: A compact convolutional neural network for EEG-based brain–computer interfaces. Journal of Neural Engineering, 15(5), 056013.

[17] Lu, Y., et al. (2024). EEGMamba: Bidirectional state space model with mixture of experts for EEG multi-task classification. arXiv preprint arXiv:2407.20254.

[18] Rodrigues, P. L. C., Jutten, C., & Congedo, M. (2019). Riemannian Procrustes analysis: Transfer learning for brain–computer interfaces. IEEE Transactions on Biomedical Engineering, 66(8), 2390–2401.

[19] Schirrmeister, R. T., Springenberg, J. T., Fiederer, L. D. J., Glasstetter, M., Eggensperger, K., Tangermann, M., et al. (2017). Deep learning with convolutional neural networks for EEG decoding and visualization. Human Brain Mapping, 38(11), 5391–5420.

[20] Wu, D., Jiang, X., & Peng, R. (2022). Transfer learning for motor imagery based brain– computer interfaces: A tutorial. Neural Networks, 153, 235–253.

[21] Wu, D., Xu, Y., & Lu, B.-L. (2022). Transfer learning for EEG-based brain–computer interfaces: A review of progress made since 2016. IEEE Transactions on Cognitive and Developmental Systems, 14(1), 4–19.

[22] Zanini, P., Congedo, M., Jutten, C., Said, S., & Berthoumieu, Y. (2018). Transfer learning: A Riemannian geometry framework with applications to brain–computer interfaces. IEEE Transactions on Biomedical Engineering, 65(5), 1107–1116.

